# Inferring Within-Host Bottleneck Size: A Bayesian Approach

**DOI:** 10.1101/116194

**Authors:** R. Dybowski, O. Restif, D.J. Price, P. Mastroeni

**Affiliations:** Department of Veterinary Medicine, University of Cambridge, Madingley Road, Cambridge, CB3 0ES, UK

**Keywords:** Bottlenecks, Bayesian inference, Wildtype isogenic tagged strains (WITS), Salmonella

## Abstract

A number of approaches exist for bottleneck-size estimation with respect to within-host bacterial infections; however, some are more appropriate than others under certain circumstances. A Bayesian comparison of several approaches is made in terms of the availability of isogenic multitype bacteria (e.g., WITS), knowledge of post-bottleneck dynamics, and the suitability of dilution with monotype bacteria. The results are summarised by a guiding flowchart.

A sampling approach to bottleneck-size estimation is also introduced.

## 1 Introduction

The outcome of an infection processes is usually underlain by a fine balance between the virulence mechanisms of the pathogen and the resistance of the host. The presence of physical, immunological or therapeutic barriers poses constraints to the ability of bacteria to divide and disseminate within the host organism. Suppose that we have a population 𝒫_0_ of bacteria. When a subset of 𝒫_0_ is inactivated by an antibiotic or an immune response, or subject to an anatomical barrier to transmission, this can result in a substantially smaller population 𝒫_1_ (commonly known as a *bottleneck*) and, after the occurrence of the bottleneck, the bacteria of 𝒫_1_ can grow to form a new population 𝒫_2_.

Understanding the site, nature and size of bottlenecks in infectious disease processes is important to rationally design prevention strategies and treatments to control the spread of the infection within a given host. In fact, classes of vaccines and therapeutic compounds differ significantly in how they restrain an infection process with respect to the control, for example, of microbial killing or division rates and the spread within and between organs.

Bottlenecks have been the focus of a number of articles, and Abel et al. (2015) provide a biologically motivated introduction to bottlenecks. Specific experimental studies that have shown bottlenecks using isogenic tagged strains include Grant et al. (2008) in the early stage of salmonellosis in mice, Schwartz et al. (2011) in urinary tract *Escherichia coli* infection in mice, Lowe et al. (2013) during *Bacillus anthracis* colonisation in mice, Kaiser et al. (2014) with *Salmonella* Typhimurium crossing the intestinal barrier in mice, Lim et al. (2014)also with *Salmonella*, Gerlini et al. (2014) and Kono et al. (2016) with invasive *Streptococcus pneumoniae*, and Abel et al. (2015) with *Vibrio cholerae* in the intestinal tract. However, in spite of the number of studies that have been made involving bottlenecks, there has not been a unified study of the various analytical methods used for tagged/multitype experimental studies, which is the motivation of this study.

## 2 Monotype populations

### 2.1 Posterior bottleneck distributions

Of interest is estimating the size of a bottleneck given observations made after the bottleneck, and possibly also before it, in a Bayesian framework. This can be expressed as the posterior probabilities

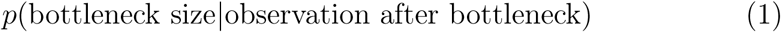

and

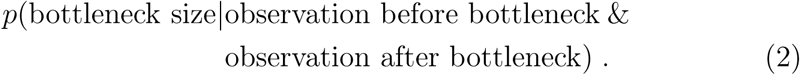

The benefit of taking the Bayesian approach of using posterior probability distributions is that such distributions not only give estimates for the most probable bottleneck sizes (in terms of the modes of the distribution) but they also express the uncertainty through the variance of the distributions.

Abel et al. (2015) estimated bottleneck size with respect to multitype populations by equating it to the effective population size as estimated by Krimbas and Tsakas (1971), which uses the standardised covariance of allele frequency due to a bottleneck. Although Abel et al. found this approach successful in the context of the within-host dynamics of *Vibro cholera*, both Pamilo and Varvio-Aho (1980) and Sourdis and Krimbas (1980) caution that bottleneck-size estimation by the Krimbas-Tsakas method can be unreliable unless sample size is sufficiently large.

Let *n*_1_ be the size of a bottleneck 𝒫_1_ and *n*_2_ the size of a post-bottleneck population 𝒫_2_ after it (Figure 1). For the posterior distribution *p*(*n*_1_|*n*_2_), Bayes’ theorem gives

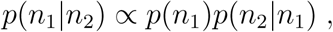

**Figure 1:**
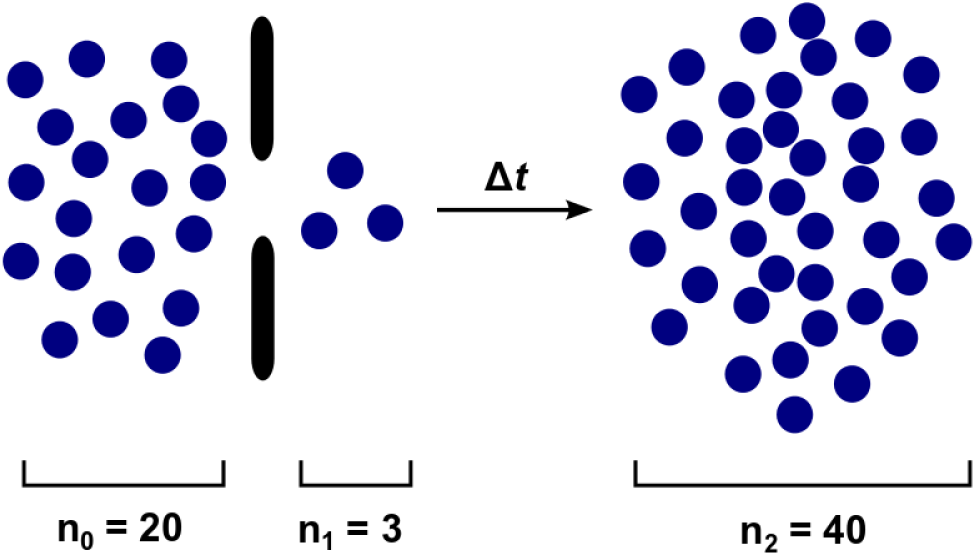
Diagrammatic representation of a bottleneck. The bottleneck is derived from a pre-bottleneck population by random sampling without replacement. A post-bottleneck population arises after time Δ*t* according to a defined growth mechanism. *n*_1_ is the size of the bottleneck, *n*_0_ the size of the pre-bottleneck population, and *n*_2_ the size of the post-bottleneck population.

and if we assume that the priors *p*(*n*_1_) are equally likely we then have the expression

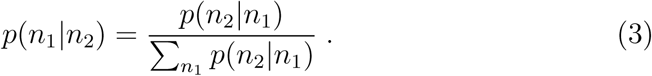

Consider *p*(*n*_2_|*n*_1_) for (3). Suppose that we can simulate the occurrence of n_2_ resulting stochastically from *n*_1_ after time interval Δ*t* (Figure 5) according to a set of parameters *θ* for the dynamics; for example, in the context of the birth-death-migration process shown in Figure 2 (Grant et al., 2008; Kaiser et al., 2014; Coward et al., 2014; Dybowski et al., 2015). Such a simulation can be achieved by a Gillespie stochastic simulation algorithm (Gillespie, 1997). If we obtain a finite number of such simulations,

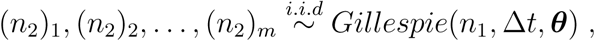

**Figure 2:**
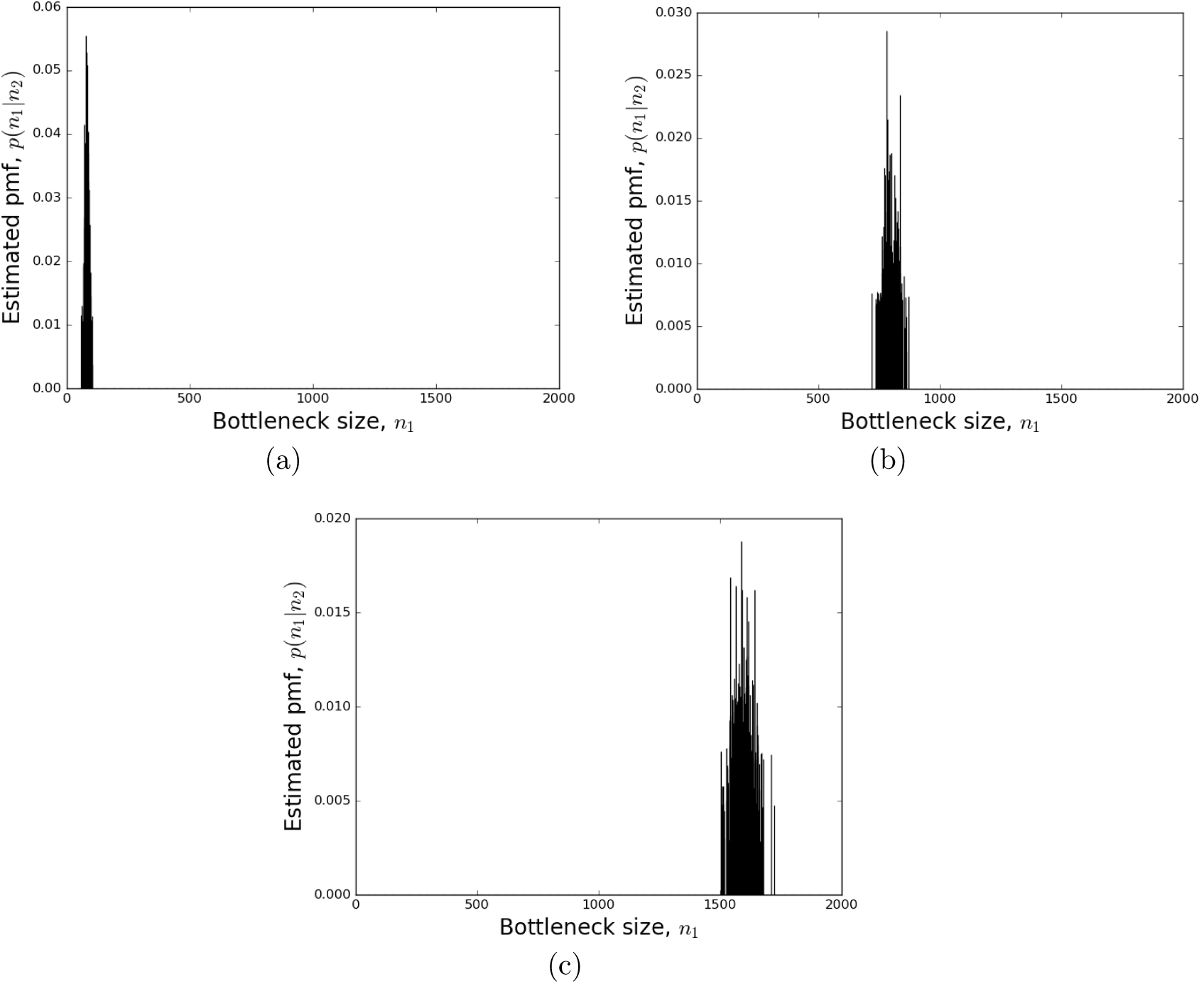
Posterior bottleneck-size distributions *p*(*n*_1_|*n*_2_) estimated using Algorithm 1. Bottlenecks of size (a) *n*_1_ = 80, (b) *ni* = 800, and (c) *n*_1_ = 1600, were artificially induced.

**Figure 3:**
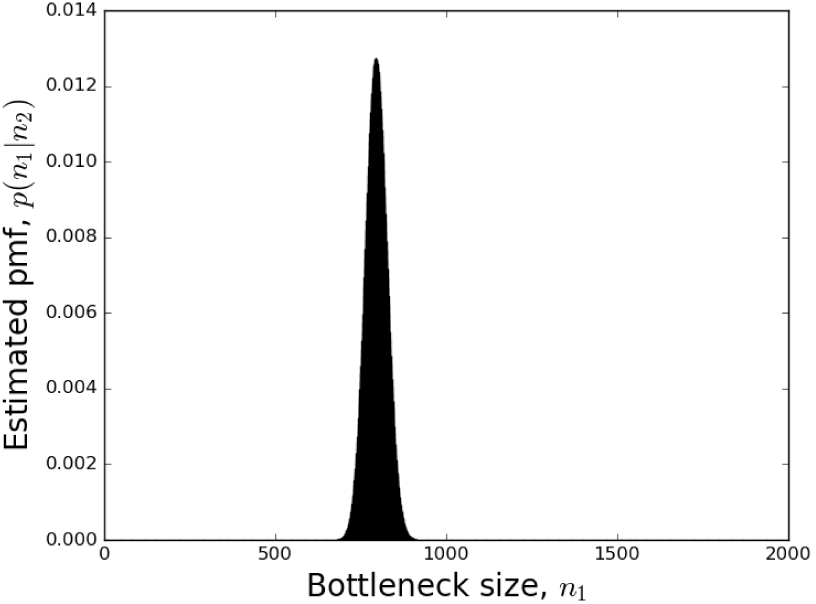
Posterior bottleneck-size distribution estimated using convolution of eight posterior distributions associated with the eight WITS. Target *n*_1_ = 800. Summary statistics are mode 793, mean 793.8, median 795 and variance 980.1. See Section 3 for details.

our task is then to estimate the probability mass function *p*(*n*_2_|*n*_1_,*θ*) from {(*n*_2_)_1_, (*n*_2_)_2_,…, (*n*_2_)_*m*_}, which can then be used for (3).

Estimating p(*n*_2_|⋅,⋅) from the sparse sample {(*n*_2_)_1_, (*n*_2_)_2_,…, (*n*_2_)_*m*_} can be attempted using local polynomial smoothing (Simonoff, 1996). A recent development in this area is the local polynomial smoothing proposed by Jacob and Oliveira (2011), which is effective for small sample sizes.

Let *ω* = {(*n*_2_)_1_, (*n*_2_)_2_,…, (*n*_2_)_*m*_}, and let the values of *ω* be placed in *k* successive cells *C*_1_,…, *C_k_*, with all occurrences of min(*ω*) being placed in cell *C*_1_, all occurrences of min(*ω*) + 1 in cell *C*_2_, …, and all occurrences of max(*ω*) in *C_k_*. The aim is to estimate the true cell probability *π_l_* for each cell *C_l_* based on the finite observations *ω*. A straightforward estimator of π_*l*_ is, of course, the relative frequency *N_l_/m*, where *N_l_* is the number of values occupying cell *C_l_*; however, using local polynomials of degree *d*, a more accurate estimation is provided by Jacob and Oliveira (2011):

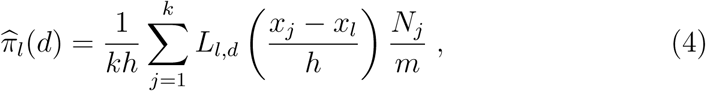

where *L_l,d_*(⋅) is the local *d*-degree polynomial estimator for the probability of cell *C_l_*, and *x_j_* = (*j* − 1/2)/*k* for *j* = 1,…, *k*. Based on the work by Ruppert and Wand (1994)on locally weighted regression, Aerts et al. (1997a, 1997b) express *L_l,d_*(.) by

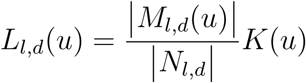

with *N_l,d_* the (*d* +1) × (*d* + 1)-matrix having the (*r*, *s*) entry given by

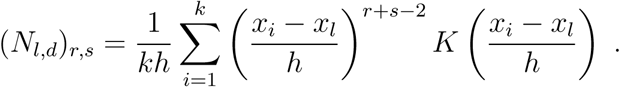

Matrix *M_l_*_,*d*_(*u*) is the same as *N_l,d_* but with the first column replaced by (1, *u*,*…*, *u^d^*)^*T*^. The width of density function *K*(.) is controlled by *h.*

Both this technique for local polynomial smoothing and Gillespie simulation were used in Algorithm 1 for the estimation of *p*(*n*_1_|*n*_2_).

#### Algorithm 1 Estimation of *p*(*n*_1_|*n*_2_).

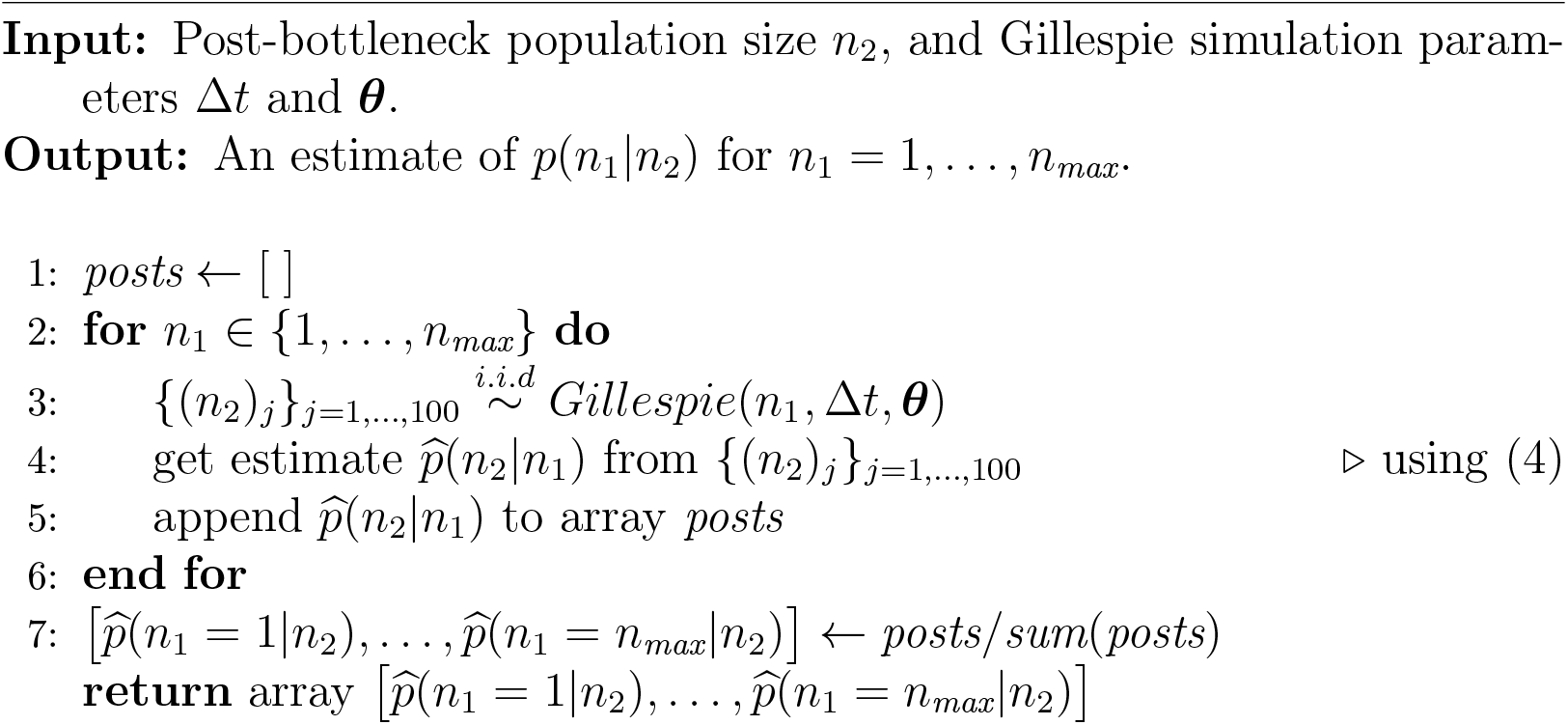

### 2.2 Example

*Salmonella enterica* is the main cause of salmonellosis, and its pathogenesis is under extensive research (Dougan & Baker, 2014). When *S. enterica* enters the bloodstream of a host, bacteria spread to a number of organs including the liver and the spleen (Figure 4). In researching this process using a mouse model, Grant et al. (2008) estimated the division, death, clearance and immigration rates of the bacterium at different times.

**Figure 4:**
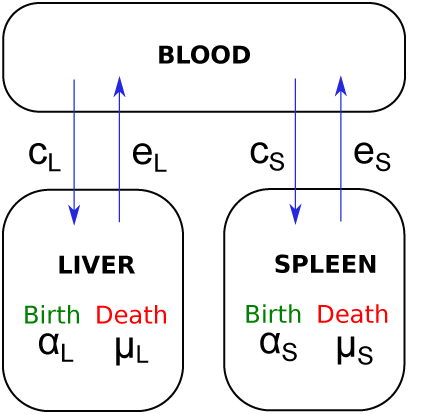
Schematic representation of the spread of *S. enterica* from the blood and between the liver and the spleen. The dynamics is governed by a set of parameters *θ*; namely, the per capita division rates (*α_L_*, α_*S*_), death rates (*μ*_L_,*μ_S_*), immigration rates (*c_L_*, *c_S_*) and clearance rates (*e_L_*, *e_S_*).

To demonstrate the efficacy of Algorithm 1 to detect bottlenecks of size 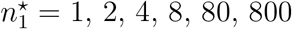 (caused by a hypothetical antibiotic treatment) in an organ such as the liver of a mouse, a value 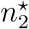 for *n*_2_ was first derived from 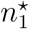 and then the posterior distribution 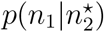 was estimated from 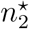 using Algorithm 1 as follows:

1. Choose a target bottleneck size 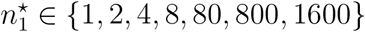
2. 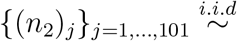 *Gillespie*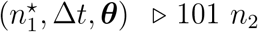 values derived from 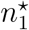
3. Set 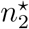 to the median of **{(***n*_2_)_*j*_**}**_*j*=1,…,101_
4. Obtain 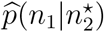 using Algorithm 1
5. Compare 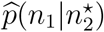 with target 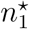

The 101 Gillespie simulations were conducted using an inoculum of 1000 bacteria at *t* = 0 (equivalent to using 101 mice). An infection was allowed to progress according to ***θ*** using the parameter values published by Grant et al. (2008) (i.e., per capita division, death, emigration and immigration rates), but at *t* = 24 hours, the number of bacteria in the liver was changed to *n*_1_ within the Gillespie simulation. The growth period Δ*t* was set to 12 hours. Estimates of p(*n*_2_**|***n*_1_,***θ***) for (3) were provided by (4) using local polynomial smoothing with an Epanechnikov kernel density function and local polynomials of degree *d* =1.

Figure 2 presents the resulting estimated posterior probabilities, and Table 1 shows that the medians of the posteriors agreed within 1% of the true bottleneck sizes.

**Table 1:**
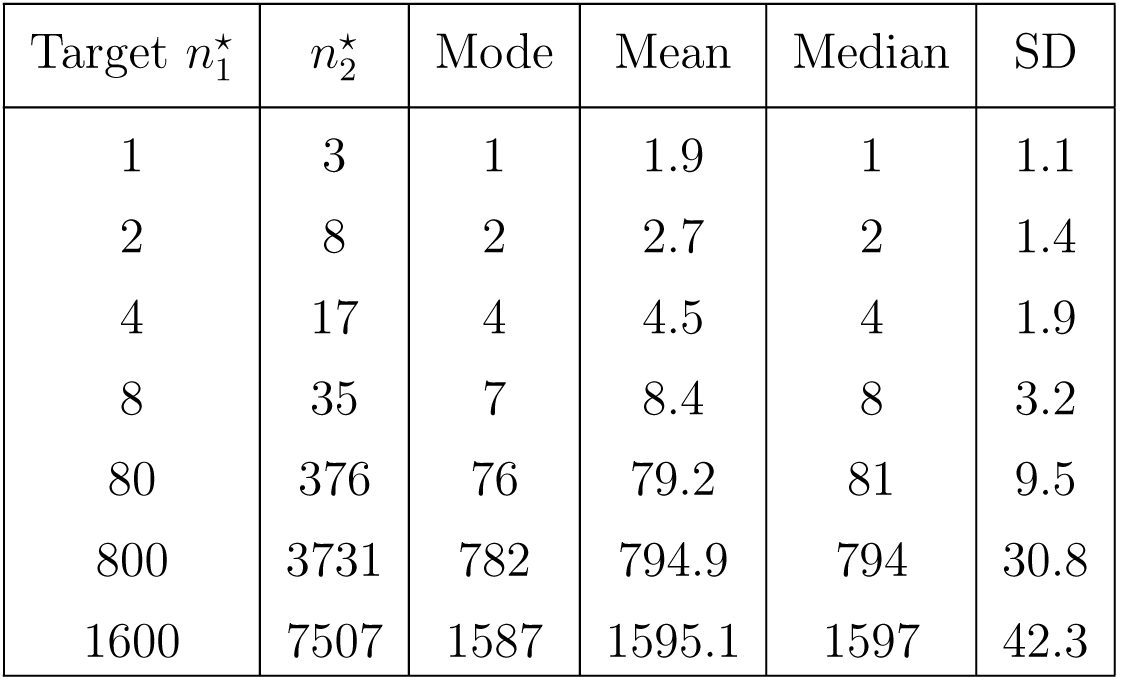
The accuracy of Algorithm 1 in terms of summary statistics for the estimated posterior distribution 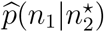. See Section 2.2 for details.

### 2.3 Inclusion of a pre-bottleneck population

What if the size *n*_0_ of the pre-bottleneck population 𝒫_0_ is also available to us, or is assumed? The posterior distribution for *n*_1_ becomes

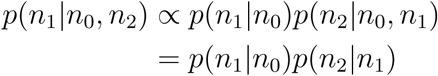

Bacteria are either killed by an antibiotic or not; therefore, with regard to *p*(*n*_1_|*n*_0_), a simple assumption is that this probability is given by the binomial probability distribution

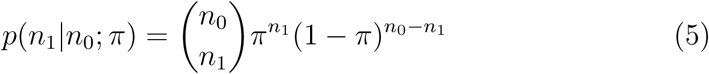

where *π* is the probability that a bacterium will be included in the bottleneck; however, *π* is not known *a priori.* Furthermore, (5) implies that the expected size *n*_1_ of the bottleneck is a linear function of *n*_0_ for all *n*_0_,

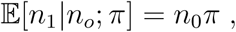

but Abel et al. (2015) warn that alternative scenarios could exist.

## 3 Multitype populations

A multitype population of bacteria is possible by using phenotypically identical bacterial strains where each strain carries a different DNA signature tag in the same noncoding region of the chromosome (Crimmins & Isberg, 2012). An example of this is the use of wild-type isogenic tagged strains (WITS) (Grant et al., 2008) which we will consider herein.

Suppose now that bacterial pre-bottleneck population 𝒫_0_ is composed of eight WITS 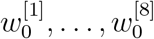, where 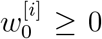 is the number of bacteria tagged with the *i***-**^th^ WITS tag. The population is reduced to bottleneck 𝒫**_1_**with WITS distribution 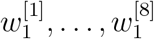, and population 𝒫_2_ resulting from the growth of 𝒫_1_ has WITS distribution 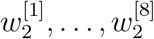. As shown in Figure 5, the distribution of WITS in 𝒫_2_ can be very different to that present in 𝒫_0_ because of the stochastic variation of 𝒫_1_ (Abel et al., 2015), and possibly also from 𝒫_1_ to 𝒫_2_.

**Figure 5:**
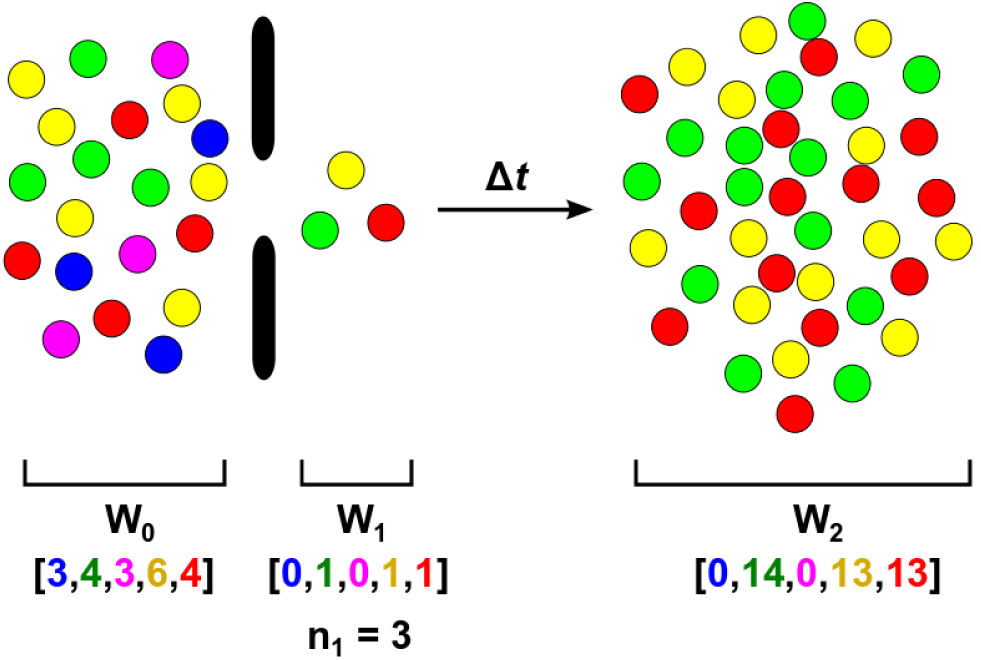
Diagrammatic representation of a bottleneck and its propensity to stochastic variability in terms of both its size (*n*_1_) and the composition of its population (**w**_1_). The array of numbers below each population states the number of tagged bacteria present in that population.

Given the WITS distribution 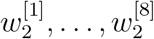 of a post-bottleneck population, estimated posterior distributions 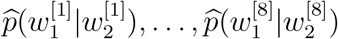 can be obtained for each of the WITS independently of each other using Algorithm 1, with 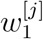 being used in the algorithm place of *n*_**1**_, and 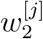 in place of *n*_2_.

The total size *n*_1_ of a bottleneck composed of WITS is given by the sum 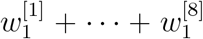. There are two approaches to estimating *n*_**1**_: (a) use the sum 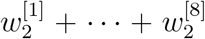] for *n*_2_, ignore the WITS tags and estimate *p*(*n*_1_|*n*_2_) using Algorithm 1; (b) determine the posterior mass function for the sum 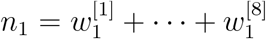 by applying convolution successively to the individual WITS posterior distributions 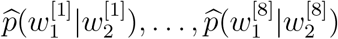.

If *X*_1_ and *X*_2_ are two independent integer-valued random variables with distribution functions *p***_1_**= *p*(*X*_1_) and *p*_2_ = *p*(*X*_2_), and *Z = X***_1_**+ *X*_2_, the distribution function *p*(*Z*) is given by

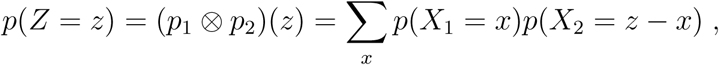

where ⊗ is the convolution operator.

Because the convolution operator is commutative, we can extend its use to sums of more than two random variables, *Z* = *X*_1_ + *X*_2_ + ⋅ + *X_m_*, by repeatedly applying the operator:

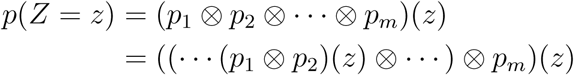

To examine the effect of using convolution, we used target value 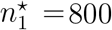 with 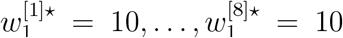. Convolution was applied to the estimated posteriors 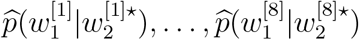 via Fourier transformation, but the resulting estimated posterior 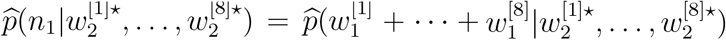 was no better than the posterior 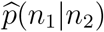 obtained by ignoring the WITS other than being smoother (Fig 6).

### 3.1 Inclusion of a multitype pre-bottleneck population

Suppose that we know, or are able to assume, the composition **w**_0_ of a prebottleneck multitype population as well as that of a post-bottleneck population. In this case, the posterior of the bottleneck size is *p*(*n*_1_|**w**_0_, **w**_2_).

Now,

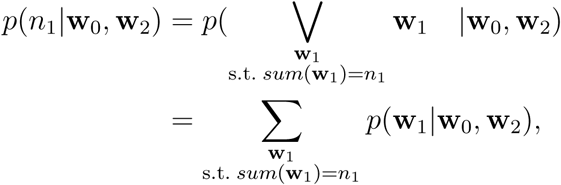

where 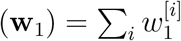. From Bayes’ theorem,

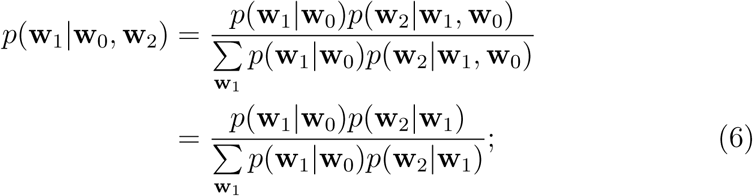

thus

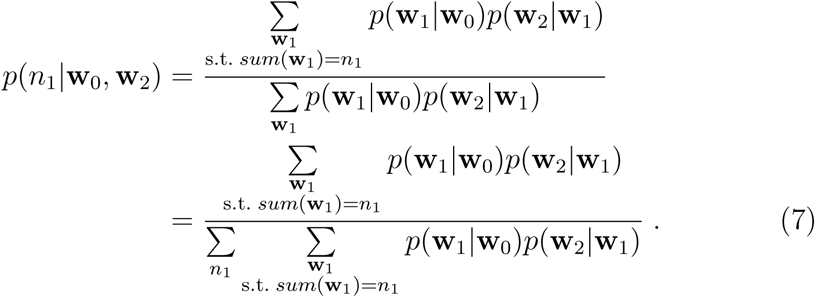

If the WITS are phenotypically identical, each bacterium has the same probability of surviving an antibiotic. Under this assumption, a distribution **w**_1_ of *n*_1_ WITS in a bottleneck 𝒫_1_ can be regarded as a sample resulting from a random selection (without replacement) of *n*_1_ WITS from the prebottleneck population 𝒫_0_ with distribution **w**_0_. The probability of selecting **w**_1_ from **w**_0_ without replacement such that *sum*(**w**_1_) = *n*_1_ is given by the multivariate hypergeometric probability distribution:

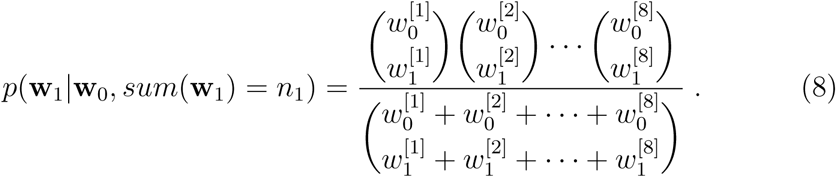

As for *p*(**w**_2_|**w**_1_), the independence between the WITS enables us to factorise *p*(**w**_2_|**w**_1_, ***θ***) as follows:

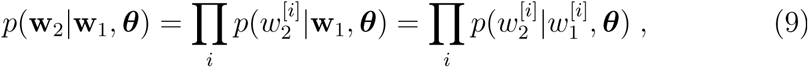

and estimation of 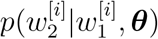 for (9) can be performed in the same manner as described for *p*(*n*_2_|*n*_1_,***θ***) using Algorithm 1.

### 3.2 On assuming proportionality

Through Bayes’ theorem, we have

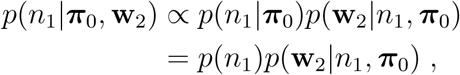

where the elements of *π*_0_ are those of **w**_0_ expressed as relative frequencies. Moreover, if we assume *p*(*n*_1_) to be equiprobable for all *n*_1_ then

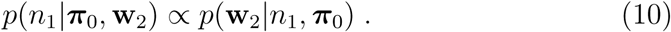

Suppose we make the further assumption that the elements of **w**_2_ developed proportionaly from those in **w**_1_: **w**_2_ = *ψ***w**_1_ for some positive integer *ψ*. This assumption implies that *ψn*_1_ = *sum*(**w**_2_) and, as *sum*(**w**_2_) is constant for a given **w**_2_, it follows that **w**_2_ can be derived from **w**_1_ using a range of *ψ* values such that *ψ* = *sum*(*w*_2_)*/n*_1_. But is one *ψ* more likely than another?

If *p*(**w**_2_\*n*_1_, *π*_0_) is defined by a multinomial distribution then

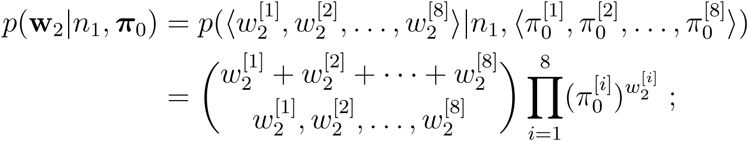

however,

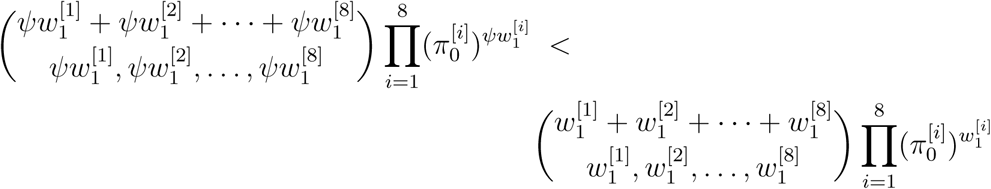

for any positive integer *ψ* (note that 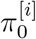 is the same on both sides of the inequality), thus

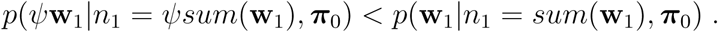

Consequently, if **w**_2_ = *ψ***w**_1_ then *sum*(**w**_1_) is the most probable value for *n*_i_. Put another way, if proportionality is assumed then the most probable value for *n*_1_ is *sum*(**w**_2_) divided by the highest common factor for the elements of **w**_2_.

### 3.3 A sampling approach

In the previous section, we have shown how to obtain values for *p*(**w**_1_**|w**_0_) and *p*(**w**_2_**|w**_1_) that are required for (7), but (7) also requires us to determine the summands for every possible **w**_1_ such that 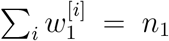. The problem with this is that the number of possible **w**_1_ for a given value of *n*_1_ grows super-exponentially with *n*_1_ (Charalambides, 2002, p.138); for example, the number of possible **w**_1_ when selecting 100 microbes from **w**_0_ = (1000^[1]^, 1000^[2]^,…, 1000^[8]^) is more than 26 thousand million. As this is combinatorially (and thus computationally) challenging, an alternative approach is required.

To circumvent the combinatorial issue, one could consider restricting the summations of (7) to the more probable configurations of **w**_1_, such as the modes of **w**_1_. An algorithm for the generation of all the modes of a multivariate hypergeometric distribution has been proposed by Requena and Cludad (2003), but a simpler approach is to randomly sample points **w**_1_, say 1000 times, from *p*(**w**_1_**|w**_0_) as a multivariate hypergeometric distribution, given that most of these points would be expected to be in the vicinity of the modes.

Our implementation of the sampling approach is shown in Algorithm 2, and its efficacy was tested using the following steps of a toy experiment:

1. Set **w**_0_ to **⟨**600^[1]^, 600^[2]^,…, 600^[8]^⟩
2. Choose a target bottleneck size 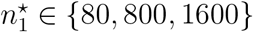
3. In order to choose a **w**_1_ associated with target 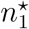, select a mode 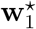 from the multivariate hypergeometric distribution associated with random samples of size 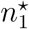 taken from **w**_0_ (Requena & Cludad, 2003)
4. In order to choose a **w**_2_ resulting from **w**_2_, first do

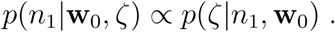
5. then, for **w**_2_, set 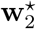 to the median of {(**w**_2_)_*j*_}_*j*=1,…,101_
6. Obtain 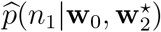 using Algorithm 2
7. Compare 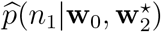 with target 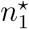

Figure 6 displays the resulting posterior distributions, which have median accuracies similar to those shown for the estimation of *p*(*n*_1_|*n*_2_) derived by Algorithm 1.

**Figure 6:**
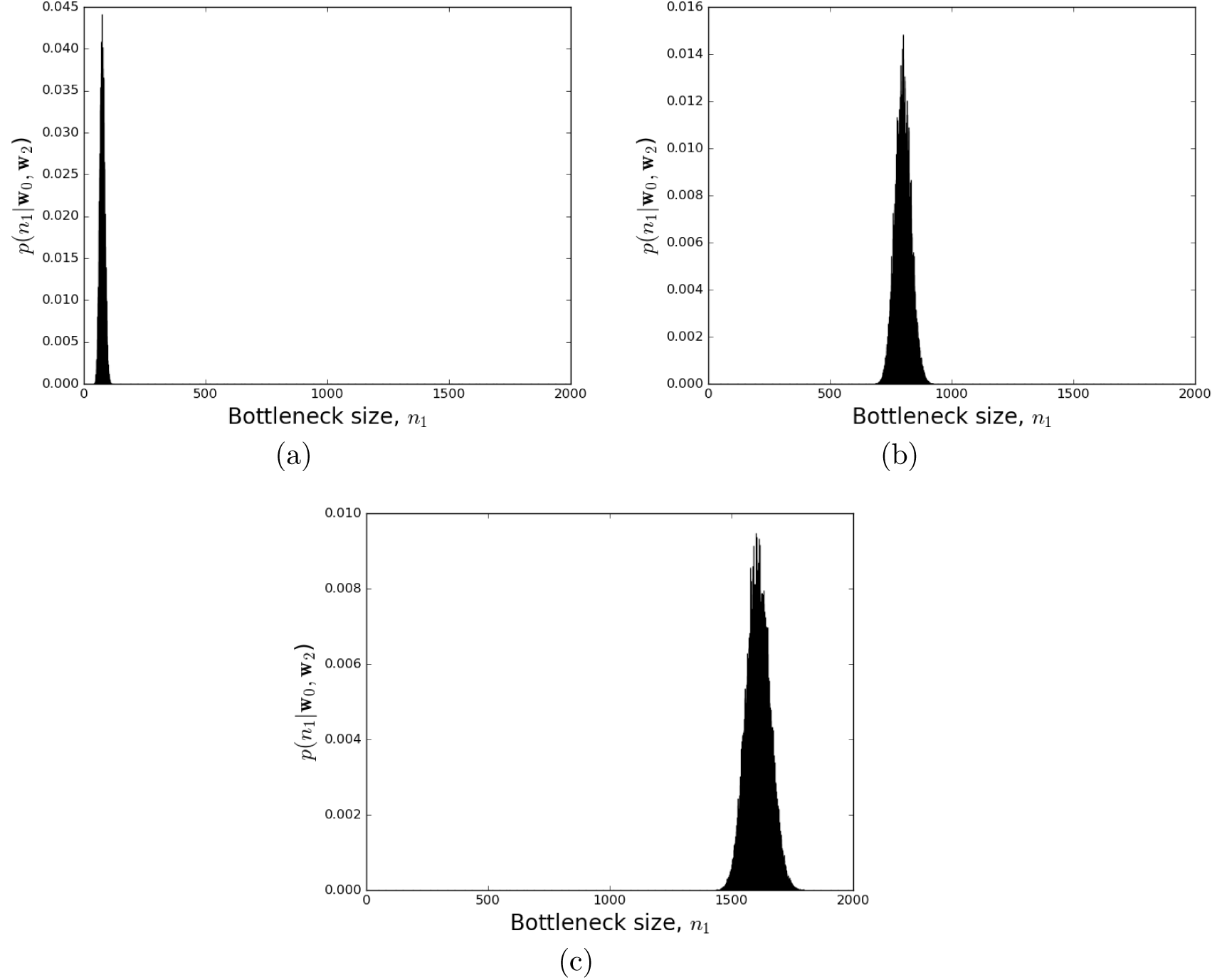
Posterior bottleneck-size distributions *p*(*n*_1_|**w**_0_, **w**_2_) estimated using Algorithm 2. Target bottlenecks of size (a) *n*_1_ = 80, (b) *n*_2_ = 800, and (c) *n*_1_ *=* 1600, were artificially induced. See Section 3.3 for details.

**Figure 7:**
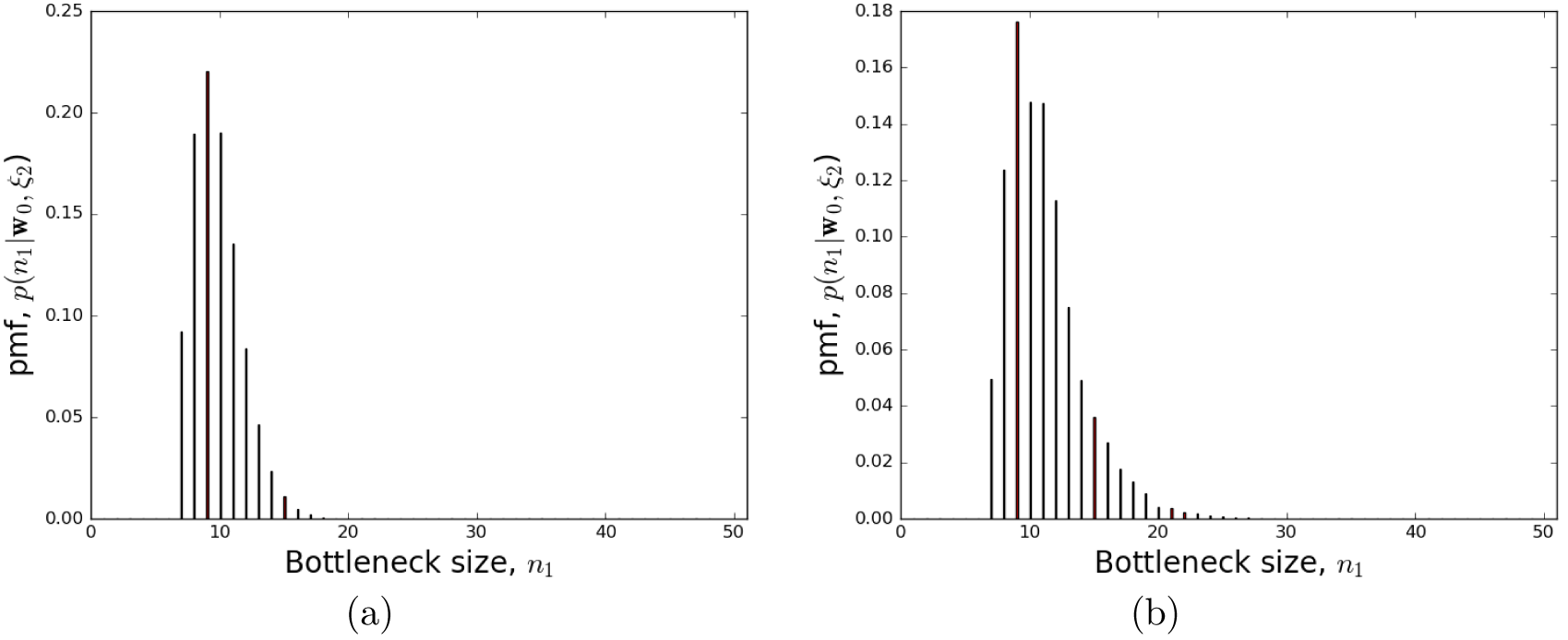
Posterior bottleneck-size distributions obtained (a) exactly using equation (12) with **w**_0_ = ⟨4^[1]^, 4^[2]^,…, 4^[8]^⟩, and (b) estimated using 1000 samples with **w**_0_ = ⟨600^[1]^, 600^[2]^,…, 600^[8]^⟩. Target is *n*_1_ = 7 in both cases.

## 4 Patterns of missing WITS

An assumption made when estimating *p*(*n*_2_|*n*_1_, ***θ***) via Gillespie simulation is that parameters ***θ*** are known and are not influenced by the presence of an antibiotic, but this is not necessarily always the case (Kaiser et al., 2014). Consequently, how can we estimate bottleneck size when ***θ*** is not known to us?

Consider (7) written as

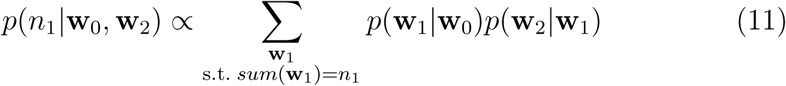

and suppose we replace **w**_2_ with a vector ξ_2_ denoting which WITS in **w**_2_ are missing, then *p*(**w**_2_|**w**_1_), in turn, becomes replaced by *p*(ξ_2_|**w**_1_). Furthermore, if we assume that missingness pattern ξ_2_ is equal to the missingness pattern ξ_1_ of **w**_1_ then ξ_2_ is implied by **w**_1_ and there is no need to consider post-bottleneck dynamics. Using this approach, (11) simplifies to

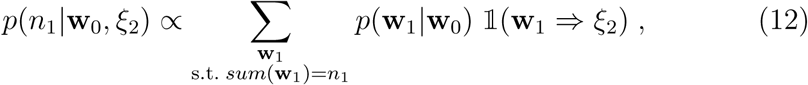

where 𝟙(⋅) is the indicator function. However, the assumption that ξ_2_ = ξ_1_, and thus that **w**_1_ ⇒ ξ_2_, may not hold if a few WITS are randomly lost soon after a bottleneck due to a combination of (a) a small bottleneck, (b) a large number of WITS, and (c) high post-bottleneck replication and death rates.

In order to compare the accuracy of (11) with (12), an experiment was used based on the following scenario. A bottleneck WITS population **w**_1_ of size *n*_1_ is assumed to have been sampled from **w**_1_ = ⟨4^[1]^, 4^[2]^,…, 4^[8]^⟩ without replacement. Each element 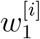 of **w**_1_ then gives rise to an element 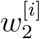 of **w**_2_ by sampling from a Poisson distribution with Poisson parameter 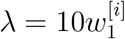. This scenario is the basis for the following toy experiment:

1. Set **w**_0_ to ⟨4^[1]^,4^[2]^,…,4^[8]^⟩
2. Choose a target bottleneck size 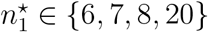
3. In order to choose a **w**_1_ associated with target 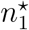, select a mode 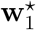 from the multivariate hypergeometric distribution associated with random samples of size 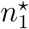 taken from **w**_0_
4. For **w**_2_, use 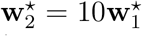 (i.e., vector of expected values from the Poisson distributions)
5. Get missingness pattern ξ_2_ corresponding to 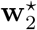
6. Obtain 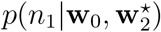 using (11)
7. Compare 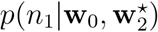 with target 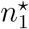
8. Obtain *p*(*n*_1_**|w**_0_, ξ_2_) using (12)
9. Compare *p*(*n*_1_**|w**_0_, ξ_2_) with target 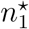

The results of the experiment are shown in Figures 8 and 9. In the case of the probability mass functions for 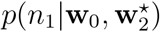), the modes coincided exactly with the target values, but this was not the case for *p*(*n*_1_**|w**_0_, ξ_2_). When at least one WITS was missing, the mode for *p*(*n*_1_**|w**_0_, ξ_2_) was greater than the target value. This is associated with the observation that the mean value of *p*(**w**_2_**|w**_1_) as encountered in (11) tended to be less than the mean value for 𝟙(**w**_1_ ⇒ ξ_2_) in (12), which is equal to *p*(**w**_1_ ⇒ ξ_2_). When no WITS were missing, the resulting probability mass function for *p*(*n*_1_**|w**_0_,ξ_2_) exhibited a plateau as *n*_1_ increased. This can be explained as follows: it is increasingly unlikely that no WITS are missing as *n*_1_ decreases; on the other hand, the absence of missing WITS can be explained by the occurrence of large *n*_1_ values up to and including the complete absence of a bottleneck.

**Figure 8:**
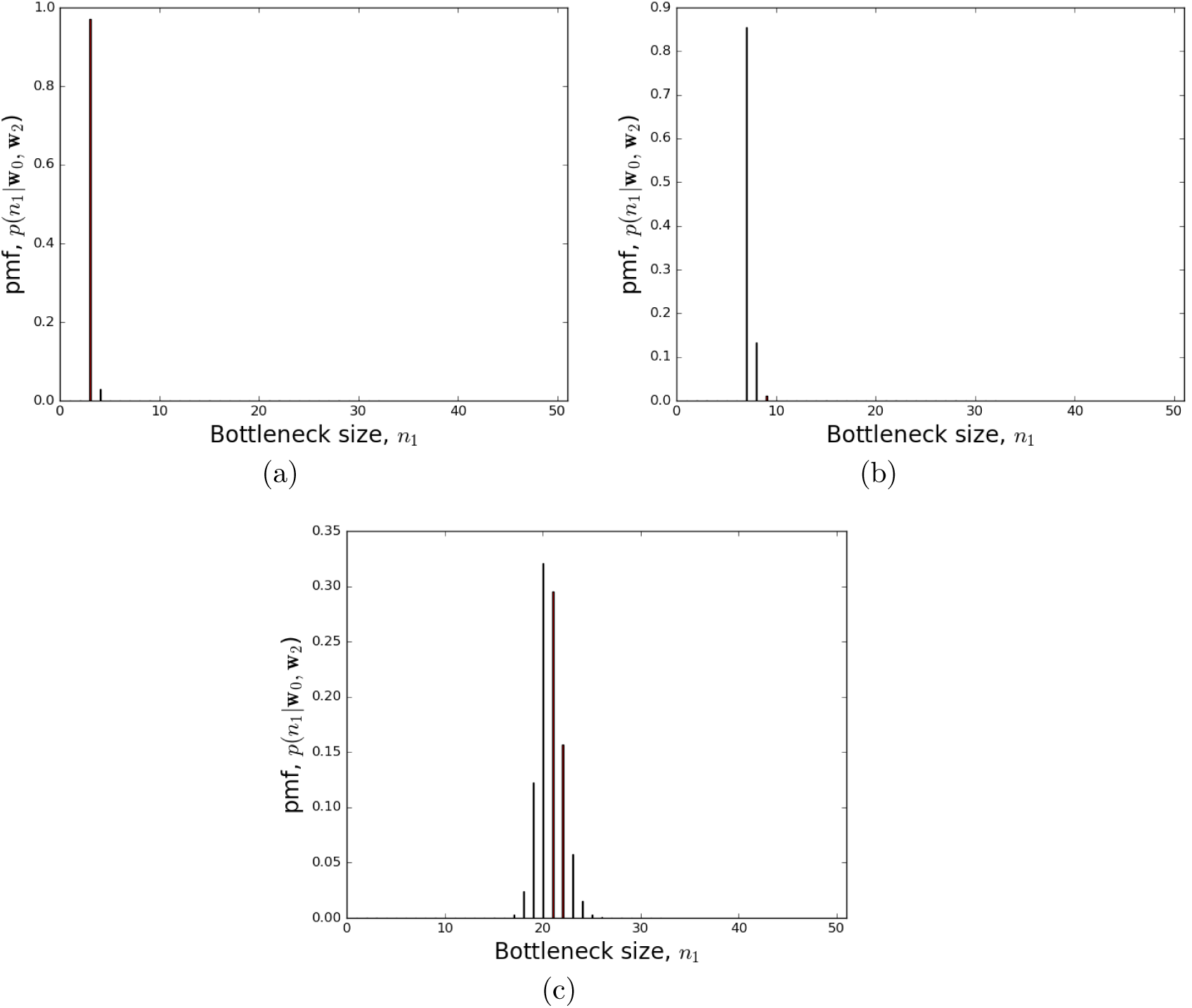
Figure 8: Posterior bottleneck-size distributions *p*(*n*_1_**|w**_0_, **w**_2_) determined accurately using (11). Target bottlenecks of size (a) *n*_1_ = 3, (b) *n*_1_ = 7, and (c) *n*_1_ *=* 20, were used. See Section 4 for details.

When *p*(*n*_1_**|w**_0_, **w**_2_) has a plateau, a lower bound for *n*_1_ can be set equal to the lower bound of the 95% highest density interval with respect to *p*(*n*_1_**|w**_0_, **w**_2_), which is a type of 95% credible interval.

Note that the above toy experiment uses Equation (12) exactly so as to display the resulting distributions. In reality, the inoculum size would be far greater than 4 × 8 and, in such circumstances, the sampling approach of Section 3.3 would be used instead. Figure shows the result of using sampling when ξ_2_ is used in place of **w**_2_, with *n*_1_ = 600 × 8.

### 4.1 Dilution of WITS with monotypes

In Figure 9 (c), the bottleneck of size 20 could not be estimated because of the presence of a plateau instead of a mode. This issue can be overcome by diluting the WITS with untagged (i.e., monotype) isogenic bacteria. The justification for this is that, for a fixed *n*_1_, the probability of at least one WITS being missing increases as the pre-bottleneck population **w**_0_ becomes more dilute.

**Figure 9:**
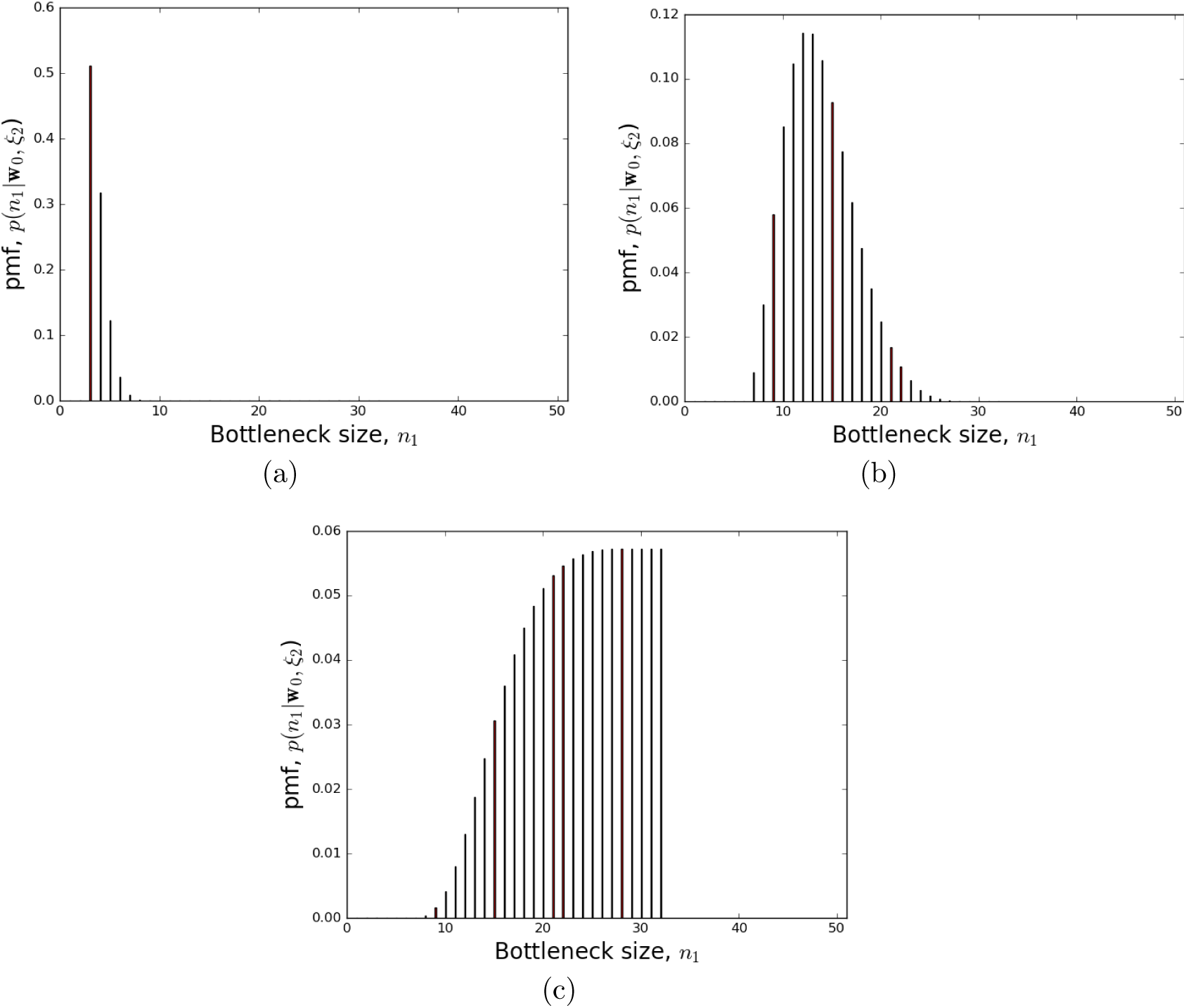
Posterior bottleneck-size distributions *p*(*n*_1_|**w**_0_,ξ_2_) determined accurately using (12). Target bottlenecks of size (a) *n*_1_ = 3, (b) *n*_1_ = 7, and (c) *n*_1_ *=* 20, were used. See Section 4 for details.

Let 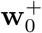 represent **w**_0_ augmented with *u* untagged bacteria. For example, if we add 10 untagged bacteria to **w**_0_ = ⟨4^[1]^,…, 4^[8]^⟩ then 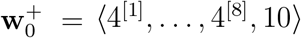 = ⟨4^[1]^,…, 4^[8]^, 10⟩. Upon using 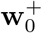 in place of **w**_0_, expression (12) becomes

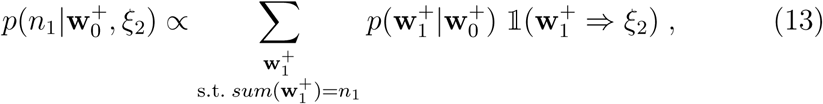

where 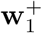 allows for the possibility that untagged bacteria can be present in the bottleneck. Implication 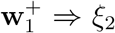 ⇒ ξ_2_ in (13) is based only on the WITS component of 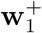; the untagged bacteria in 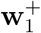 are ignored.

By way of example, suppose that we add *u* = 13 untagged bacteria to **w**_0_ = ⟨4^[1]^,…, 4^[8]^⟩ in order to perform the following toy experiment:

1. Set 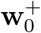 to ⟨4^[1]^, 4^[2]^,…, 4^[8]^,*u*⟩
2. Set number of untagged bacteria *u* =13
3. Set target bottleneck size 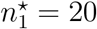
4. In order to choose a 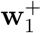 associated with target 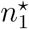, first select a mode 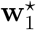 from the multivariate hypergeometric distribution associated with random samples of size 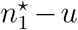 taken from ⟨4^[1]^, 4^[2]^,…, 4^[8]^⟩, and then set 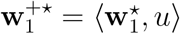
5. For **w**_2_, use 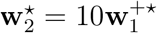
6. Get missingness pattern ξ_2_ corresponding to 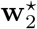
7. Obtain 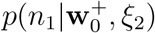 using (13)

Figure 10 shows that dilution with untagged bacteria has allowed an estimate of the bottleneck size to be performed. But a note of caution is due. Using *u* =13 allowed the bottleneck size to be estimated as 18 (whereas *u* =12 gave a plateau), but the position of the mode for 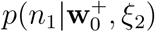 is influenced by the choice of *u*, with the mode decreasing as *u* increases. For example, the mode was 13 when *u* =14 and 10 when *u* = 15. A similar behaviour has been observed when using other target values for *n*_1_. This suggests that one should use the smallest possible value for *u* that permits a mode to appear instead of a plateau.

**Figure 10:**
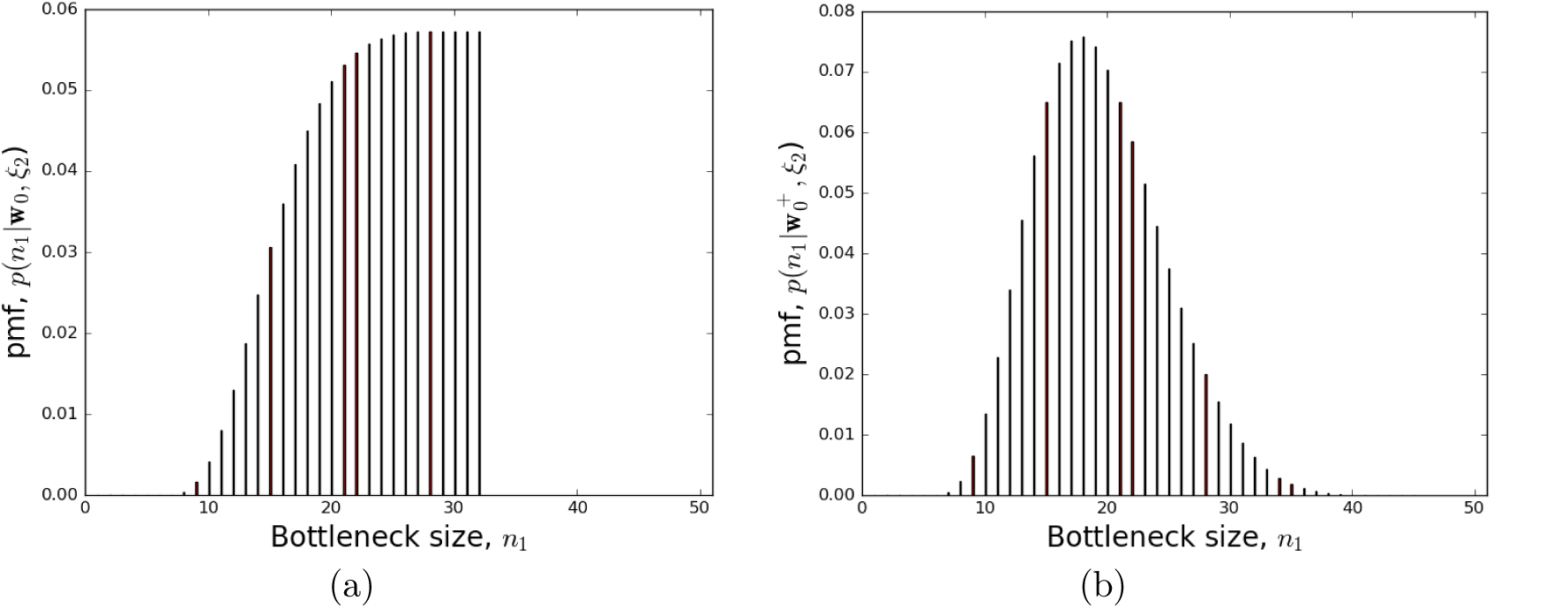
Posterior bottleneck-size distributions (a) without dilution and (b) with dilution. Distributions *p*(n_1_|**w**_0_,ξ_2_) and 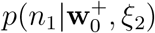 were determined accurately using (12) and (13), respectively. The target bottleneck size was *n*_1_ = 20 in both cases. For the dilution, *u* = 13 untagged bacteria were present in 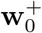. See Section 4.1 for details.

The above WITS dilution technique was used by Maier et al. (2014) to estimate the size of gut luminal bottlenecks during *Salmonella* Typhimurium colitis. The inoculum consisted of seven WITS in equal proportions, which was increasingly diluted with an untagged isogenic wild-type strain until a loss of at least one WITS was first detected in a post-bottleneck population. This point occurred at a dilution of 1:7000. The size of a bottleneck was then estimated using a likelihood function based on binomial selection.

Lim et al. (2014) also developed a method to estimate the size of a bottleneck from an observation of missing WITS, but their approach was more granular in that it was restricted to just those cases where at least one WITS was missing but without considering the precise the number of missing WITS. Let ζ ∈ {*true*, *false*} denote the state that at least one WITS is missing from **w**_2_ (and thus assumably from **w**_1_). In this context, the posterior for bottleneck size is *p*(*n*_1_|**w**_0_, ζ). The computational approach used by Lim et al. differed from (12) in that they derived a plot of 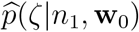 as a function of *n*_1_ using computer simulations and then assumed that

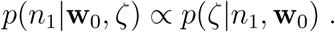

### 4.2 On increasing the number of WITS

All the examples shown so far have been based on the use of eight WITS, but what if a larger number of WITS are used? Intuitively, if we keep the bottleneck size *n*_1_ constant but increase the number **|***𝒲***|** of WITS available then the number of missing WITS is expected to increase. Furthermore, the rate of change in the number of missing WITS as *n*_1_ decreases is expected to increase as **|***𝒲***|** increases. This suggests that accuracy in the estimation of *n*_1_ from missingness patterns *ξ*_2_ should improve with larger **|***𝒲***|**. This argument is supported by the results of the computer simulations conducted by Lim et al. (2014)using different values for **|***𝒲***|** in which a significant improvement occurs on going from **|***𝒲***|** = 10 to **|***𝒲***|** = 40.

As a further demonstration, the posterior distribution *p*(*n*_1_**|w**_0_,ξ_2_) shown in Figure 9 when *n*_1_ = 7, which is based on 8 WITS, was recalculated using 12 WITS resulting in a decrease in variance (Figure 11).

**Figure 11:**
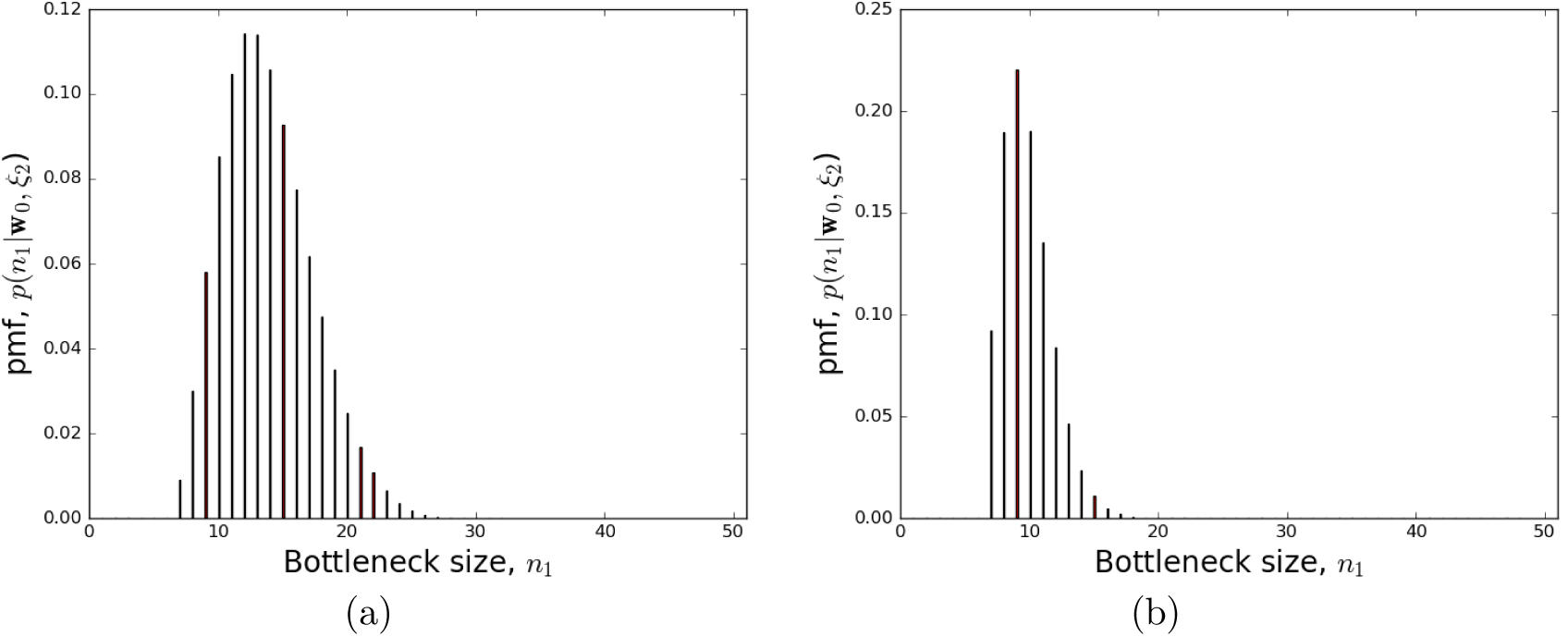
Posterior bottleneck-size distributions obtained (a) with 8 WITS and (b) with 12 WITS. Note the decrease in variance.

## 5 Discussion

We describe Bayesian approaches to estimating the size of bacterial bottlenecks given observation of a post-bottleneck population (either monotype or multitype), but size estimation with respect to monotypes is possible only when post-bottleneck dynamics is known (or assumed).

The use of multitype isogenic bacteria in the form of WITS allow the composition of pre-bottleneck populations to be included in the analysis. Furthermore, the use of WITS enables bottleneck sizes to be estimate when post-bottleneck dynamics is not known. This is done through observation of patterns of missing WITS; however, this can require dilution of an inoculum with isogenic untagged bacteria. Figure 12 provides a flowchart that summarises these observations.

**Figure 12:**
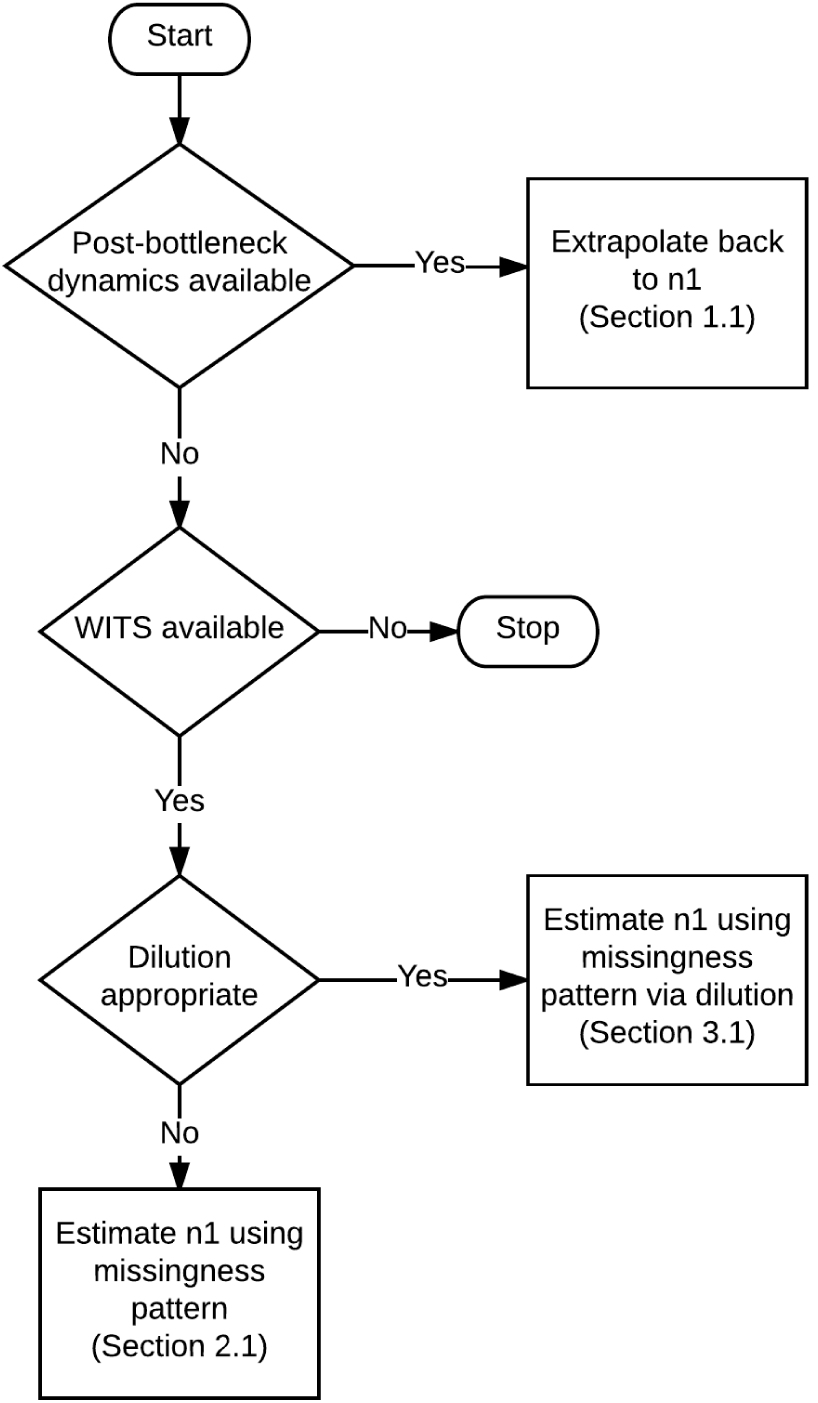
Flowchart summarizing alternative strategies to bottleneck-size estimation depending on circumstances.

## Author contributions

RD, PM and OR conceived the project. RD performed the mathematical analyses and simulations, which were checked by DJP. All the authors contributed to the preparation of the manuscript.

## Acknowledgements

RD was supported by BBSRC grant BB/I002189/1 awarded to PM. DJP was supported by BBSRC grant BB/M020193/1 awarded to OR.

## Algorithm 2 Estimation of *p*(*n*_1_|**w**_0_, **w**_2_).

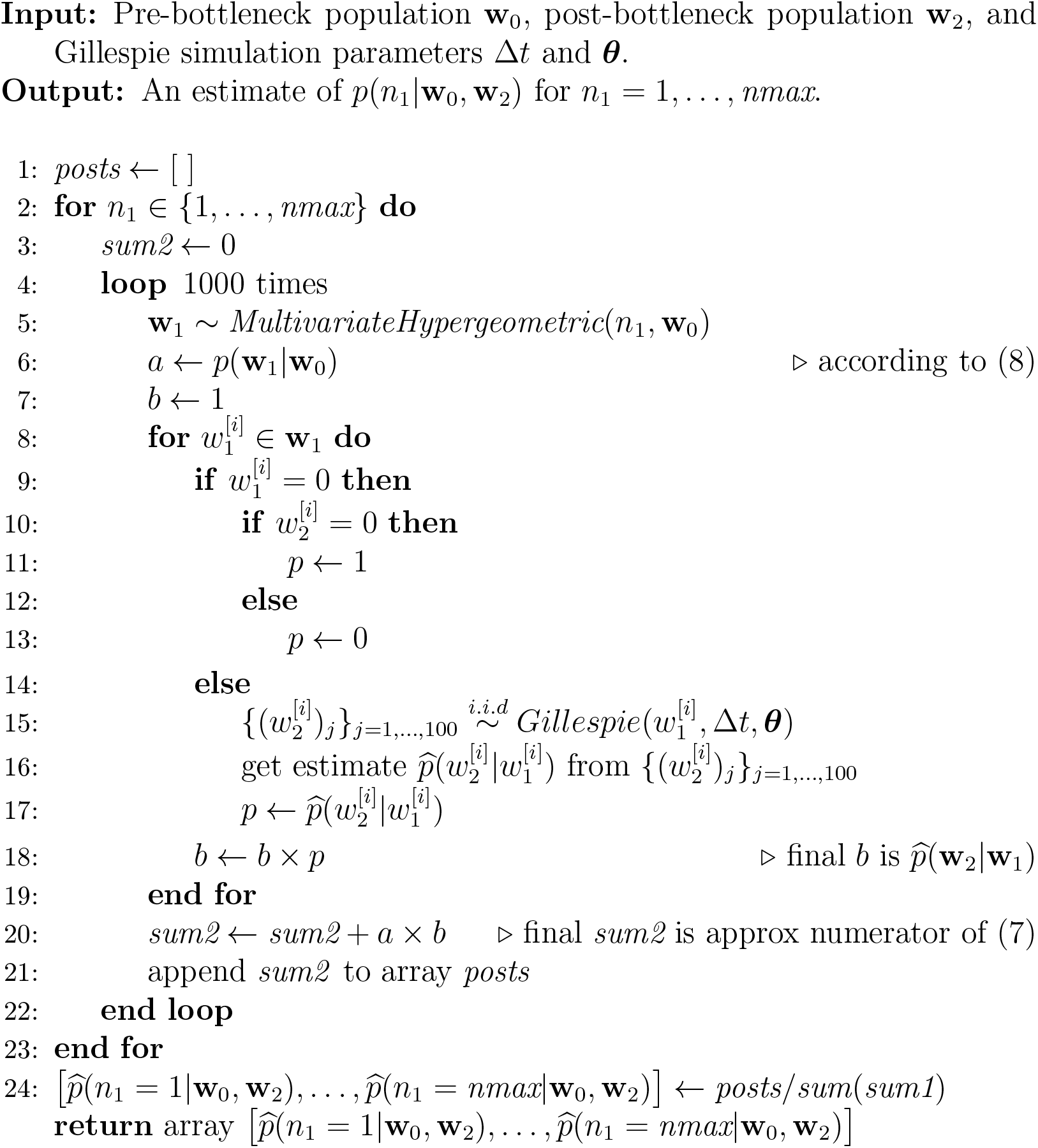

